# Panmixia across elevation in thermally sensitive Andean dung beetles

**DOI:** 10.1101/783233

**Authors:** Ethan Linck, Jorge E. Celi, Kimberly S. Sheldon

**Affiliations:** Department of Ecology & Evolutionary Biology, University of Tennessee, Knoxville; Biogeography and Spatial Ecology Research Group, Universidad Regional Amazónica Ikiam

**Keywords:** climatic variability hypothesis, mountain passes, isolation-by-environment, Scarabaeinae, tropics, Ecuador

## Abstract

Janzen’s seasonality hypothesis predicts that organisms inhabiting environments with limited climatic variability will evolve a reduced thermal tolerance breadth compared with organisms experiencing greater climatic variability. In turn, narrow tolerance breadth may select against dispersal across strong temperature gradients, such as those found across elevation. This can result in narrow elevational ranges and generate a pattern of isolation-by-environment, or neutral genetic differentiation correlated with environmental variables that is independent of geographic distance. We tested for signatures of isolation-by-environment across elevation using genome-wide SNP data from five species of Andean dung beetles (subfamily Scarabaeinae) with well-characterized, narrow thermal physiologies and narrow elevational distributions. Contrary to our expectations, we found no evidence of population genetic structure associated with elevation and little signal of isolation-by-environment. Further, elevational ranges for four of five species appear to be at equilibrium and show no evidence of demographic constraints at range limits. Taken together, these results suggest physiological constraints on dispersal may primarily operate outside of a stable realized niche.

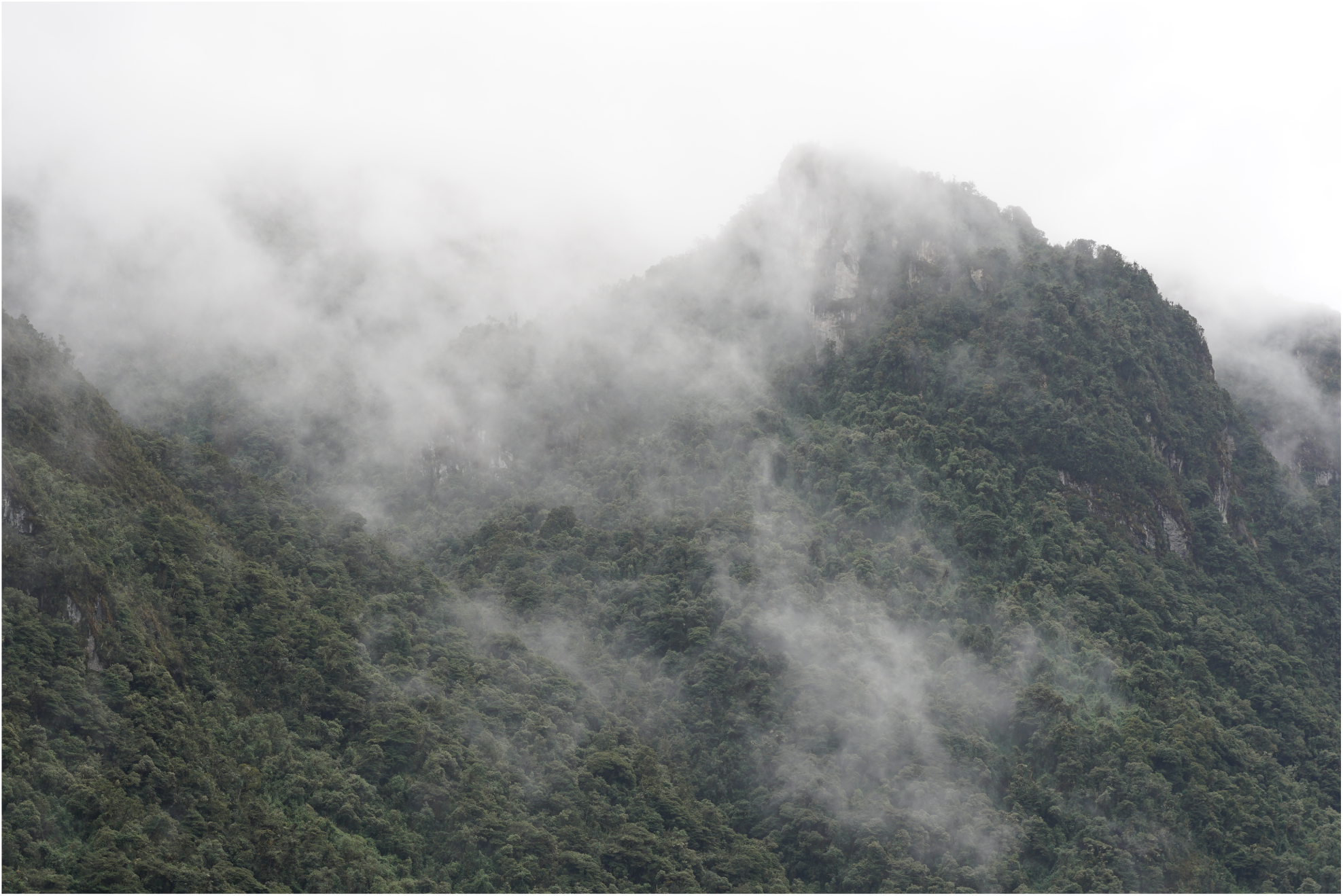

## Introduction

The movement of individual organisms has profound consequences for biogeography, ecology, and evolution. Dispersal and its absence shape range limits (Kirkpatrick & Barton, 1997; Sexton, McIntyre, Angert, & Rice, 2009), community assembly and disassembly (Cody, MacArthur, Diamond, & Diamond, 1975; Sheldon, Yang, & Tewksbury, 2011), and a species’ ability to track its niche under a changing climate (Schloss, Núñez, & Lawler, 2012; Urban, Tewksbury, & Sheldon, 2012). When followed by interbreeding between immigrants and residents, dispersal influences rates of gene flow between physically separated populations. In turn, rates of gene flow can affect demography and population structure (Bohonak, 1999; Slatkin, 1985, 1987), probabilities of extinction (Soulé, 1987; Tallmon, Luikart, & Waples, 2004; Whiteley, Fitzpatrick, Funk, & Tallmon, 2015), adaptive potential (Aitken & Whitlock, 2013; Garant, Forde, & Hendry, 2007; García-Ramos & Kirkpatrick, 1997; Lenormand, 2002), and ultimately speciation and biogeography (Cadena et al., 2012; Kisel & Barraclough, 2010; Mallet, 2008). Yet despite its obvious importance, general predictors of dispersal rate remain elusive (Bowler & Benton, 2005; Johnson & Gaines, 1990).

A theory that is a notable exception in attempting to predict dispersal rates of organisms is Dan Janzen’s seasonality hypothesis, which mechanistically links temperature variation across latitude with its consequences for dispersal and biogeographic patterns (Janzen, 1967). The hypothesis begins with the assumptions that 1) tropical ecosystems generally have less seasonal variation in temperature than temperate ecosystems and that 2) populations and species are adapted to the range of climates they experience. As a result, Janzen predicted that a tropical organism climbing a mountainside is more likely to encounter a physiologically challenging climate than a temperate organism would be, leading to selection against dispersal across elevational gradients. A suite of downstream predictions naturally follow, e.g. that tropical organisms should also have narrower elevational ranges, show increased population subdivision and a pattern of isolation by environment, and ultimately have higher speciation rates (Ghalambor, Huey, Martin, Tewksbury, & Wang, 2006; Wang & Bradburd, 2014; Gadek et al., 2018; Sheldon, Huey, Kaspari, & Sanders, 2018).

Tests of Janzen’s seasonality hypothesis across taxa have supported its physiological assumptions and predictions, albeit with caveats (Ghalambor et al., 2006; Sheldon et al., 2018). Salamanders (Feder, 1982) and lizards (van Berkum, 1988) show narrower ranges of body temperature in the tropics, suggesting the limited thermal variability they directly experience is strongly correlated with regional climate. Dung beetles (Sheldon & Tewksbury, 2014) and montane stream insects (Polato et al., 2018), as well as amphibians (Snyder & Weathers, 1975) and insects (Abraham et al., 2000) more broadly show increased thermal tolerance (as measured by CTmax - CTmin) at higher latitudes, though this effect is reduced in the more aseasonal southern hemisphere. On the other hand, evidence for reduced thermal plasticity in tropical organisms—which Janzen expected in aseasonal environments for the same reason as reduced thermal tolerance, i.e. as its costs would outweigh its benefits—is mixed at best (Brattstrom, 1968; Feder, 1982; Tsuji, 1988; Gunderson & Stillman, 2015). Studies have generally found that elevational ranges are narrower in tropical species (Terborgh, 1977; Huey, 1978; Rahbek & Graves, 2001; Sheldon & Tewksbury, 2014; Gadek et al., 2018), though endotherms may be an exception to this pattern (Sheldon, Yang, & Tewksbury, 2011; Cadena et al., 2012).

Despite reduced thermal tolerance and generally smaller elevational ranges of tropical species, surprisingly few researchers have examined dispersal across tropical gradients, though it is the mechanism in the seasonality hypothesis that links physiology with elevational ranges. Population genetic structure across elevation in Andean sparrows (Cheviron & Brumfield, 2009) and subtropical forest trees in China (Shi, Michalski, Chen, & Durka, 2011) indicate gene flow across mountain sides can be sufficiently reduced to permit genome-wide differentiation. Evidence for speciation across elevational gradients—a possible long-term consequences of population genetic structure—has been reported in tropical kingfishers (Linck, Freeman, & Dumbacher, 2019) and butterflies (Elias et al., 2009). To our knowledge, the only paper explicitly estimating effective migration rates across elevational gradients found reduced gene flow and greater population subdivision in tropical stream insects compared to their temperate relatives (Polato et al., 2018). However, the authors did not control for the possibility that dispersal is systematically reduced in the tropics for reasons other than physiological tolerance, e.g. by comparing the relative contributions of isolation by environment and isolation by distance across latitude.

We used a densely sampled population genomic dataset to ask whether mountain passes are “higher” in five species of dung beetles (Scarabaeinae) with well-characterized thermal physiologies and elevational distributions that conform to predictions of Janzen’s seasonality hypothesis. We test for reduced dispersal across elevational gradients relative to within elevational bands by estimating fine-scale population genetic structure, Wright’s neighborhood size, and by using a Bayesian approach for describing the relative contribution of geographic distance and environmental distance to neutral genetic differentiation. To understand whether observed patterns are temporary artifacts of recent population expansion, we further ask whether elevational ranges are at equilibrium, and test for evidence of demographic constraints at elevational range limits.

## Materials and Methods

### Study system and sampling

The true dung beetles (subfamily Scarabaeinae) are increasingly popular organisms in studies of ecology and evolution (Simmons & Ridsdill-Smith, 2011; Hanski & Cambefort, 2014). Ectotherms with a global distribution, they are useful taxa for comparative studies of natural history, physiology, and population genetics. Our previous work using phylogenetically matched dung beetles from locations spanning 60° of latitude found thermal tolerance of species in the tribes Canthonini and Dichotomini generally increased with seasonality and was positively correlated with elevational range width—key predictions of Janzen’s seasonality hypothesis (Sheldon & Tewksbury, 2014). We focused our current study on two species in the tribe Canthonini (*Deltochilum speciosissimum, Deltochilum tessellatum*), two species in the tribe Dichotomini (*Dichotomius podalirius, Dichotomius satanas*) and a single species (*Eurysternus affin. flocossus*) from a third tribe, Oniticellini. Species in the genus *Deltochilum* are ball rolling dung beetles that feed and breed using dung or carrion, though at least one tropical species of *Deltochilum* is predatory and kills millipedes (Larsen et al. 2009). Beetles in the tribe Dichotomini excavate tunnels near or below dung deposits and then transport it underground to create brood balls for laying and incubating eggs, often closing off the tunnel’s entrance (Hanski & Cambefort, 2014). Finally, *Eurysternus* spp. are unique among dung beetles in several aspects of their reproductive biology, including a “nuptial feast”, or aggregation and consumption of dung balls prior to nesting, an inability to roll balls with their legs, multiple nests consisting of a shallow crater with several brood balls, nest care, and pair bond behavior in some species (Halffter, Halffter, & Huerta, 1980).

We sampled beetles using pitfall traps baited with human dung (Davis, Scholtz, & Chown, 1999) along two elevational and two horizontal transects on the eastern slope of the Andes Mountains in Napo Province, Ecuador (**Figure 1**). Our two elevational transects spanned 730 to 1175 m and 1500 to 1950 m in largely undisturbed humid forest, with four sampling localities spaced as close to 125 m apart as possible given local constraints of soil and topography. Our two horizontal transects were each approximately 2 km long, with one transect having four points and the other having two points. Both horizontal transects were located in replicate corridors of montane forest at 2150 m asl separated by a natural barrier, the Cosanga River. No species were shared between the two elevational transects, reflecting high beta diversity in Andean scarabs (Sheldon & Tewksbury, 2014). In all cases, we sampled nearly the complete elevational range of our focal species (K. Sheldon, unpublished data.) Following capture, all beetles were euthanized and stored in 95% EtOH prior to processing.

**Figure 1:**
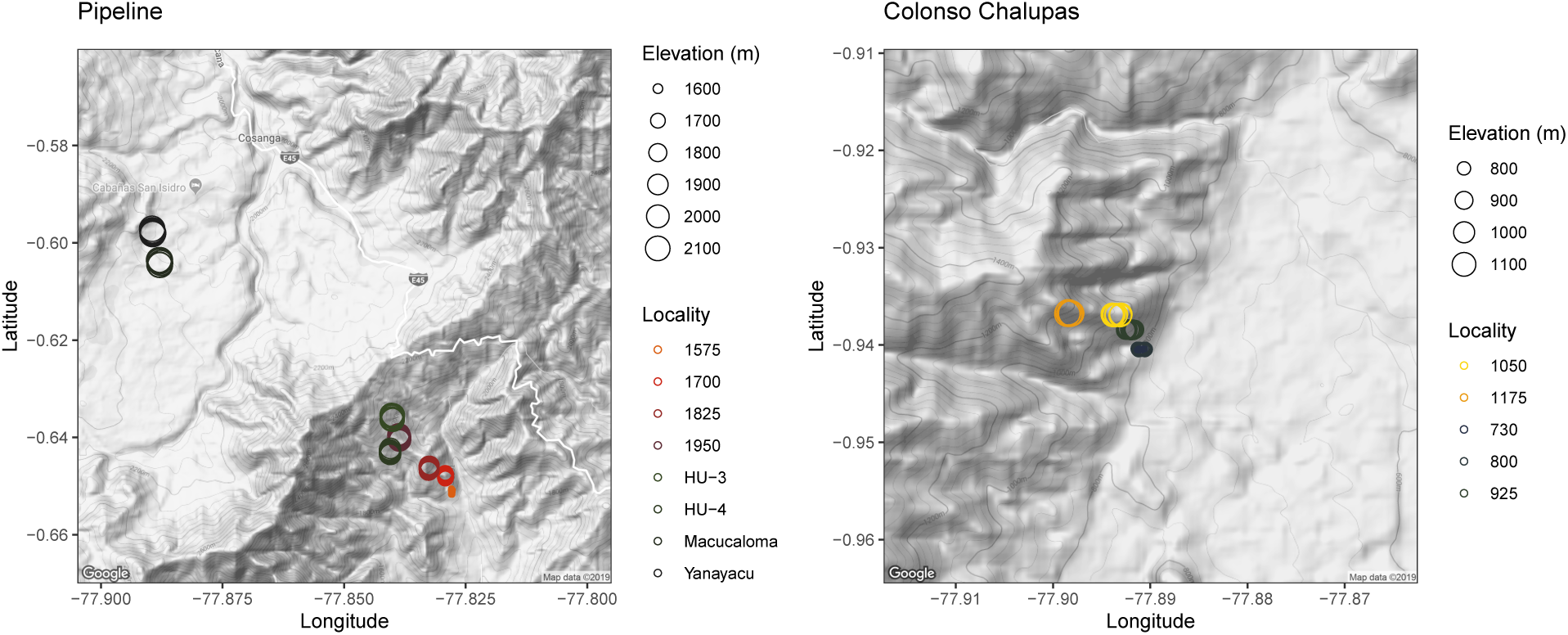
Sampling localities across two elevational transects in Napo Province, EC.

### Library preparation and DNA sequencing

We extracted genomic DNA from all samples using Qiagen DNeasy kits and the manufacturer’s recommended protocol for insects, homogenizing a small amount of wing muscle tissue in pH 7.2 PBS prior to lysis (Qiagen, Hilden, Germany). To prepare libraries for reduced representation high throughput sequencing, we used a double restriction enzyme digest approach adapted from Gompert et al., 2014 and described below.

We placed 6 *μ*L DNA from each sample (with a minimum concentration of 20 ng/ *μ*L) in separate wells of a chilled 384 well plate, filling the remainder with samples from a separate but related study. We then added 2.6 *μ*L of a restriction digest master mix consisting of 10X T4 buffer, 1M NaCL, 1 mg/mL BSA, water, and the MseI and EcoRI enzymes in a 1:0.52:0.52:0.2125:0.10:0.25 ratio. After sealing, vortexing, and centrifuging the plate, we incubated it for 2 hours at 37°C on a thermocycler with a heated lid, followed by 20 minutes at 65°C to denature the active enzymes. We next added 1.44 *μ*L of an adaptor ligation master mix to each well. This master mix consisted of a MseI-specific adaptor, water, 10x T4 buffer, 1M NaCl, 1 mg/mL BSA, and T4 DNA ligase in a 1:0.072:0.1:0.05:0.05:0.1675 ratio. We independently added 1 *μ*L of a set of uniquely barcoded EcoRI adaptors to each well, permitting pooled samples to later be identified and demultiplexed. Again sealing, vortexing and centrifuging the plate, we incubated it on a thermocycler at 16°C for 2 hours, then diluted the reaction by adding 3 *μ*L to 19.5 *μ*L water on a new plate.

To enrich our libraries, we performed two separate 20 *μ*L PCR amplifications, each of which was conducted according to the following protocol. We added 3 *μ*L of our libraries to 17 *μ*L of a PCR mix in a new plate, and performed the reaction using a thermocycler profile of 98°C for 30 seconds, 30 cycles of 98°C for 20 seconds, 60°C for 30 seconds, 72°C for 40 seconds, and final extension at 72°C for 10 minutes. Our PCR mix consisted of water, 5x iProof Buffer, dNTPs, 50mM MgCl2, 5*μ*M Illpcr1 and Illpcr2 oligos, iProof polymerase, and DMSO in a 10.4:4.0:0.4:0.4:1.3:0.2:0.3 ratio. We then performed an additional PCR cycle to eliminate any remaining single stranded DNA and reduce sequencing errors, adding 2.1 *μ*L of a new master mix consisting of 5x iProof buffer, 5 *μ*M Illpcr1 and Illpcr2 oligos, and 10 *μ*M dNTPs in a 0.425:1.3:0.4 ratio to each well. We ran this additional cycle at 98°C for 3 minutes, 60°C for 2 minutes and 72°C for 10 minutes.

After merging our replicate PCR reactions to reduce stochastic differences in sequencing effort, we selected fragments between 250 bp and 250 bp using a Blue Pippin machine (Sage Science, Beverly, MA). We confirmed fragments observed the size expected size distribution using a bioanalyzer and sent the plate for 100 bp single end read Illumina sequencing at the Genome Sequencing and Analysis Facility (GSAF) & DNA Sequencing Facility at The University of Texas (Austin, Texas).

### Sequence assembly and variant calling

After initially demultiplexing our libraries using our unique barcodes and a Python script implemented in the first step of ipyrad (Eaton, 2014), we assembled sequencing reads and called and filtered variants for each species independently using the dDocent pipeline (v. 2.7.8), a set of bash wrapper scripts optimized for population genomics of nonmodel organisms requiring de novo assemblies (Puritz, Hollenbeck, & Gold, 2014). dDocent first removes Illumina adapter sequences and low-quality reads using Trim Galore (v. 0.6.2) (http://www.bioinformatics.babraham.ac.uk/projects/trim_galore/), itself a wrapper around the CutAdapt (Martin, 2011) and FastQC (http://www.bioinformatics.babraham.ac.uk/projects/fastqc/) tools. The pipeline then assembles reads into loci within individuals using the RADseq assembly program Rainbow (Chong, Ruan, & Wu, 2012) allowing a maximum number of 6 mismatches, and aligns loci across individuals using CD-HIT (v. 4.8.1) (Fu, Niu, Zhu, Wu, & Li, 2012). CD-HIT requires a user-input similarity threshold, which we set to 0.9 for *Deltochilum speciosissimum, Deltochilum tessellatum, Dichotomius podalirius, and Eurysternus affin. flocossus*. We used a threshold of 0.8 for *Dichotomius satanas*, as exploratory data analysis suggested high genetic diversity and we had poor assembly performance with more stringent parameter values. dDocent next calls on BWA-MEM (v. 0.7.17) (Li, 2013) to align reads to the assembled reference in BAM format; here we used default parameter values. After read alignment, dDocent uses BEDtools (v. 2.28.0) to create intervals along contigs with high quality mapping scores, which are piped to FreeBayes (v. 1.3.1) (Garrison & Marth, 2012) for Bayesian variant detection, taking advantage of the total coverage for a given base across all individuals. The pipeline concatenates single nucleotide polymorphism (SNP) and insertion-deletion calls from FreeBayes into a single .vcf file using VCFtools (v. 0.1.16) (Danecek et al., 2011).

We then used VCFtools, vcffilter (1.0.0) (https://github.com/vcflib/vcflib), and the dDocent script dDocent_filters to filter this file to high quality SNPs alone. We first dropped any individuals with missing data at more than 30% of loci, any loci missing data at more than 25% of individuals, all SNPs with a minimum minor allele frequency of 0.05 and a minimum minor allele count of 3, all SNPs with a quality (Phred) score of <30, and any genotypes with fewer than 5 reads across all individuals. We subsequently dropped sites with an allelic balance of >0.75 or <0.25, where the proportion indicates the ratio of reference allele reads to all reads; sites with reads from both strands; sites with a ratio of mapping qualities between reference and alternate alleles that were >0.9 or <1.05, and sites with a quality score >¼ below its depth. Finally, we identified sites with a depth exceeding 3X the square of the mean, and removed any from this subset that lacked quality scores exceeding 2X their depth or were outliers, which we identified qualitatively based on the approximate point at which a histogram of mean depth across loci began to asymptote. For the remainder of the paper we refer to genotypes filtered in this way as our ‘primary’ SNP datasets.

### Describing population genetic structure

To estimate population genetic structure within species across our gradients, we used two approaches appropriate for large SNP datasets. First, we performed principal component analysis (PCA) of genotypes using our primary SNP datasets using adegenet v. 2.1.1. We plotted samples on principal component axes 1 and 2, and visually identified outliers representing misidentified samples, cryptic species, or highly divergent subpopulations. Second, we tested for fine-scale population subdivision within each species using fineRADstructure v. 0.3.2, which uses haplotype data to identify a closest relative for each allele and then sums these data into coancestry similarity matrix among all individuals. We again used our primary SNP datasets, but dropped individuals identified as potentially belonging to different species through our PCA.

### Testing for isolation-by-environment

We quantified the relative contributions of geographic distance and environmental distance to observed genetic differentiation within all five species using BEDASSLE v. 1.5 (Bradburd, Ralph, & Coop, 2013). BEDASSLE is a Bayesian method that models allele frequencies in unlinked loci in a set of populations as a spatially correlated Gaussian process. Covariance is a decreasing function of both ecological and geographic distance, and parameters are estimated with a Markov Chain Monte Carlo simulation. We first filtered out primary SNP datasets for linkage disequilibrium using bcftools v. 1.9 and a maximum r2 value of 0.1, and converted these files into allele counts per population (sampling locality) using adegenet v. 2.1.1 and custom R code (https://github.com/elinck/scarab_migration/blob/master/scripts/bedassle.R). We calculated geographic distances among all sampling localities using the earth.dist() function in the R package fossil (Vavrek, 2011), and calculated environmental distances using the R package raster (Hijmans et al., 2019) and data for elevation, mean annual precipitation, and mean annual temperature from WorldClim2 (Fick & Hijmans, 2017). To promote more efficient chain mixing, we standardized both distance matrices by dividing values by their standard deviations and stored standard deviation constants to later back-transform our results (G. Bradburd, pers. comm.). We tuned MCMC parameters by conducting short runs of 10,000 generations, changing step sizes for parameters until preliminary results suggested convergence on a stationary distribution and (if possible) acceptance rates fell between 20% and 70%. After discovering computational time was prohibitive for the species with the most data (*D. tessellatum*), we reduced its SNP matrix to 2000 sites through random sampling. We then ran the MCMC for 2 million generations for each species, sampling every 250 generations, and assessed convergence by examining whether parameter values and log-likelihood values approximated stationarity. After discarding the first 25% of generations as burnin, we calculated the mean and standard deviation of the posterior probability of the ratio of isolation-by-environment to isolation-by-distance using the remaining MCMC output.

### Demographic modeling

To test whether elevational ranges were in equilibrium or the product of recent population expansion, we compared simple demographic models of drift-mutation equilibrium and exponential population growth using an approximate Bayesian computation approach (Beaumont, Zhang, & Balding, 2002). We first calculated the site frequency spectrum (SFS) of a separate SNP dataset treated identically to our primary SNP dataset except for filters based on minimum minor allele count and minor allele frequency using the R package adegenet’s glSum() function (Jombart, 2008). We next defined demographic models of a single population with either a single population-scaled nucleotide diversity parameter *θ* (our “null”) model, or both *θ* and an exponential population growth parameter *α* (our “growth”) model using the coalescent simulator framework coala v. 0.5.3 in R (Staab & Metzler, 2016). Under the “growth” model, the population size changes by a factor *e*^−*αt*^, where *t* is the time in generations since the growth has started. To approximate our empirical SNP data, we used scrm to simulate 50 three-nucleotide diploid loci for a sample size equivalent to each of our five species after filtering using the sequential coalescent with recombination model (Staab, Zhu, Metzler, & Lunter, 2015). We ran 100,000 simulations for each and calculated the resulting SFS. We then used the R package abc v. 2.1 to estimate parameters and perform model selection for each species (Csilléry, François, & Blum, 2012). We first performed leave-one-out cross validation to evaluate the ability of ABC to distinguish among our models using tolerance rates of 0.01, 0.05, and 100 simulations. We then estimated parameters and performed model selection using the abc() and postpr() functions, implementing the rejection algorithm with a tolerance rate of 0.05 for both.

### Spatial patterns of genetic diversity and Wright’s neighborhood size

To describe genetic diversity across the elevational distribution of our focal species, we calculated the population-scaled nucleotide diversity of each sampling locality using an estimator of theta (*θ* = 4*N* _*e*_*μ*) based on the mean homozygosity of gene frequencies in the R package pegas v. 0.1.1. (Paradis, 2010). We used our primary SNP dataset and calculated theta values for each RAD locus independently, plotting their full distribution. We examined the relationship between theta and the absolute distance in meters from mean sampling elevation (as a proxy for proximity to putative range limits) using linear mixed effects models with population as a fixed effect and tested for statistical significance using a likelihood ratio test. We additionally calculated Wright’s neighborhood size for each species, a metric proportional to the average number of potential mates for an individual given its dispersal ability and defined as *N*_*w*_ = 4*πρσ*^2^, where *σ* is mean parent-offspring distance and *ρ* is population density (Battey, Ralph, & Kern, 2019; Wright, 1946). To do so, we used Rousset’s finding that the reciprocal of the slope of a linear regression of the natural log of geographic distance against *F*_*ST*_ / (1-*F*_*ST*_) is an estimator of 4*πρσ*^2^ (Rousset, 1997). We analyzed pairwise *F*_*ST*_ values from BEDASSLE’s calculate.all.pairwise.Fst() function and pairwise geographic distances calculated as described above (Bradburd et al., 2013). We transformed variables and ran simple linear regressions using custom R code (https://github.com/elinck/scarab_migration/blob/master/stats.R).

## Results

### Sampling and DNA sequencing

In total, we extracted DNA from 230 individuals of our five focal species: *Dichotomius satanas* (n=100), *Dichotomius podalirius* (n=26), *Dichotomius satanas* (n=100), *Dichotomius podalirius* (n=26), Deltochilum speciosissimum (n=49), *Deltochilum tessellatum* (n=20), and *Eurysternus affin. flocossus* (n=35). Coverage was largely even across species, with mean read count ranging from 824,833.5 for *Eurysternus affin. flocossus* to 917,563.1 for *D. satanas*. After assembly and quality filtering of data, we dropped 1 individual of *D. satanas*, 2 individuals of *D. speciosissimum*, 1 individual of *D. tessellatum*, and 11 individuals of *E. affin. flocossus*. Our primary SNP datasets showed marked heterogeneity in both the number of loci and the resulting number of SNPs across species, with a minimum of 73 assembled RAD loci and 147 SNPs for D. podalirius and a maximum of 27,379 loci and 54,890 SNPs for *D. tessellatum*. As exploratory tuning of assembly parameters did not qualitatively change our results, we believe this variation is an artifact of underlying template quality, relative genetic diversity, or structural variation. Our secondary SNP dataset, which was not filtered by minor allele frequency or count, also showed substantial variation (albeit with less dramatic extremes), ranging from 347 loci and 1314 SNPs for *E. affin. flocossus* to 19,068 loci and 68,373 SNPs for *D. speciosissimum*.

### Population genetic structure

Principal component analysis and fineRADstructure found no evidence of population genetic structure associated with elevation in any species. After dropping outlier samples representing putative cryptic species, the first principal component of individual genotypes explained a minimum of 3.27% of variation (in *D. speciosissimum*) and a maximum of 23.74% of variation (in D. podalirius). Sampling elevation was uncorrelated with PC1 across the 5 species (p>0.05 and R2<0.05 for all tests) (**Figure 2A**). Similarly, coancestry matrices estimated with fineRADstructure (reflecting patterns of recent coalescence) showed no clustering of individuals by elevation in any species (**Figure 2B**).

**Figure 2:**
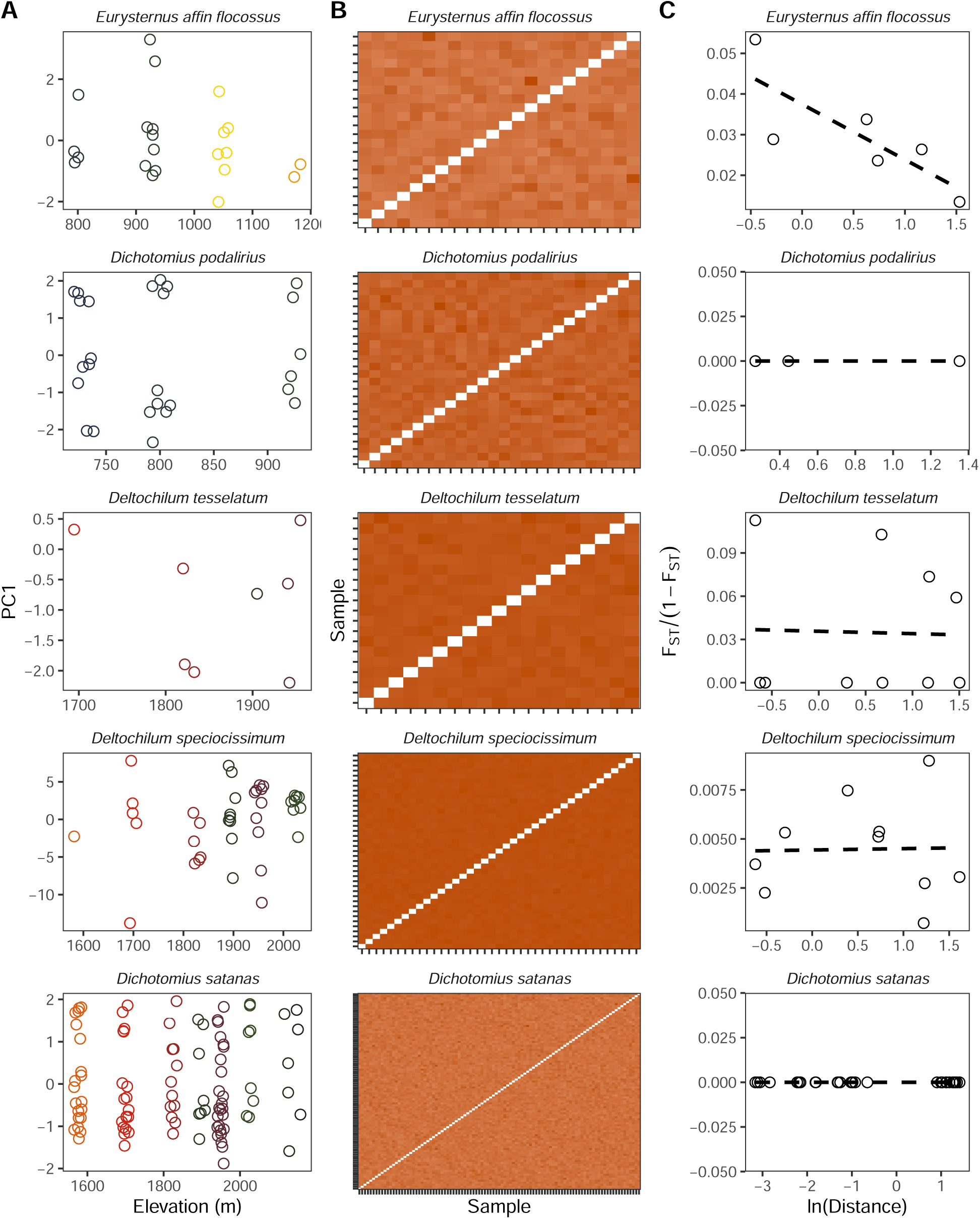
Panmixia across elevation in Andean dung beetles. A) The relationship between genotype PC1 and elevation by species. B) Coancestry matrices from pairwise estimates of individual relatedness. Rows and columns are ordered by elevation and sampling locality. Values are scaled from 0 to 1 to be comparable across species. C) The relationship between the natural log of geographic distance and standardized *F_ST_* scores. Negative or flat slopes indicate panmixia.

### Isolation-by-environment

Bayesian analysis of the relative contribution of isolation-by-environment (IBE) and isolation-by-distance (IBD) to genetic differentiation in BEDASSLE found little evidence of IBE in any focal species (**Figure 3**). Mean ratios (±SD) for the posterior probability of the parameter *α*E (reflecting the contribution of environment) to aD (reflecting the contribution of geographic distance) were 0.0006 (±.0002) for *D. satanas*, 0.0303 (±0.4417) for *D. speciosissimum*, 0.0006 (±0.0001) for *D. tessellatum*, 0.0171 (±0.0406) for D. podalirius, and 0.0221 (±0.176) for *E. affin. flocossus*. Overall, low global *F*_*ST*_ values (ranging from -0.036 for *D. satanas* to 0.029 for *D. speciosissimum*; negative values reflect higher than expected heterozygosity when using Weir and Hill’s *θ* as an estimator (Weir and Hill 2002)) likely impeded MCMC efficiency: in all but one species, acceptance rates fell to near 0 following burnin.

**Figure 3:**
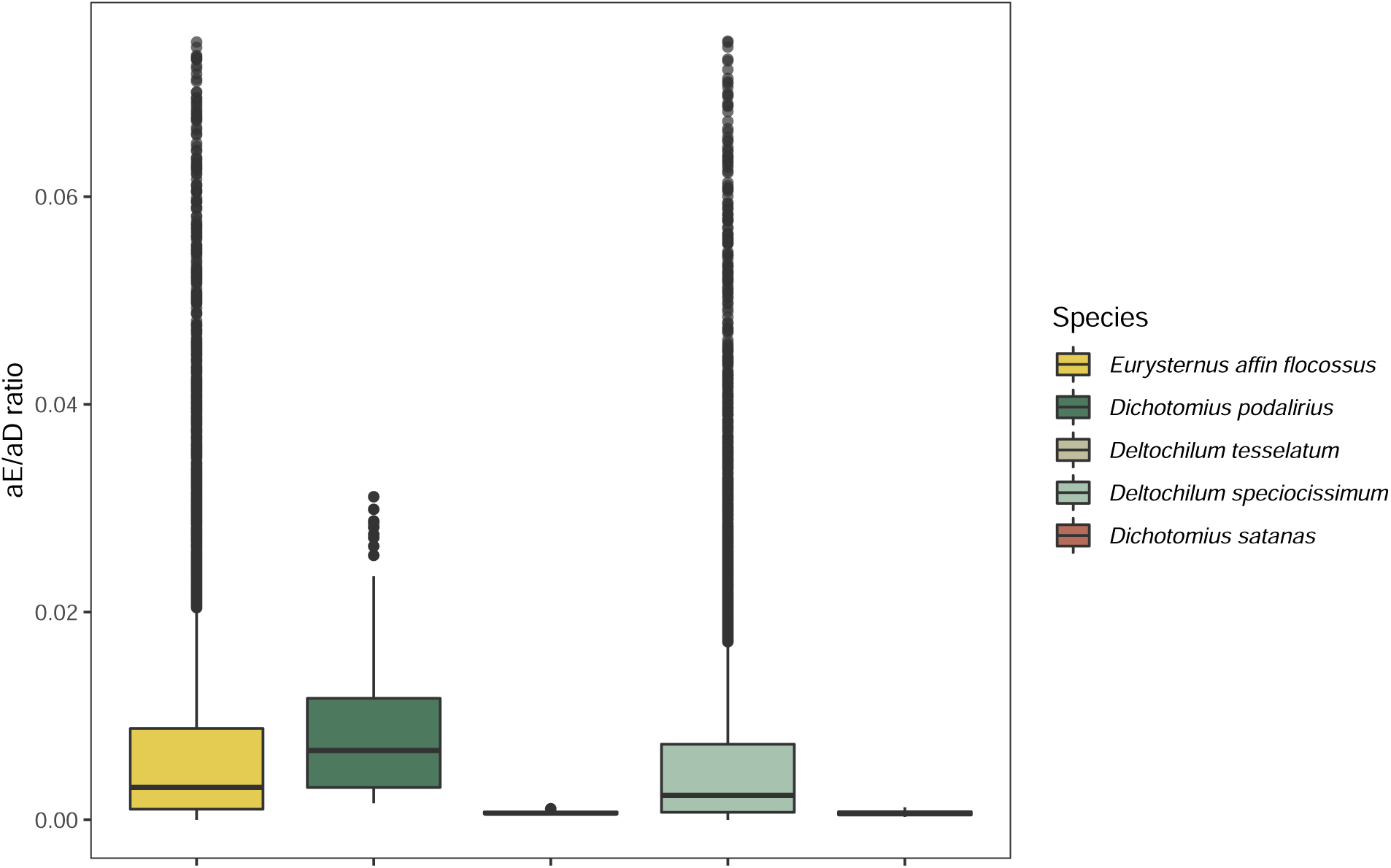
Little evidence for isolation-by-environment in Andean dung beetles. Posterior probability distributions of the ratio of the relative contribution of isolation-by-environment to isolation-by-distance from BEDASSLE. The long tail of the distributions of *E. affin. flocosusus* and *D. speciocissimum* has been trimmed to facilitate visual comparison across species.

### Demographic modeling

Demographic model testing with approximate Bayesian computation suggests four out of five focal taxa have not experienced recent range expansion (**Figure 4A**). Bayes factor values for a model representing a null hypothesis of no recent population growth compared to a model representing a hypothesis of exponential population growth were 5.165 for *D. satanas*, 3.814 for *D. speciosissimum*, 7.605 for *D. tessellatum*, and 4.081 for D. podalirius, all consistent with moderate to strong support (Kass & Raftery, 1995). In contrast, the Bayes factor value for a comparison of population growth to the null model for *E. affin. flocossus* was 1.631, indicating low to moderate support. These patterns were reflected in the posterior probabilities of the population growth rate parameter in coala (“r” in **Figure 4B**).

**Figure 4:**
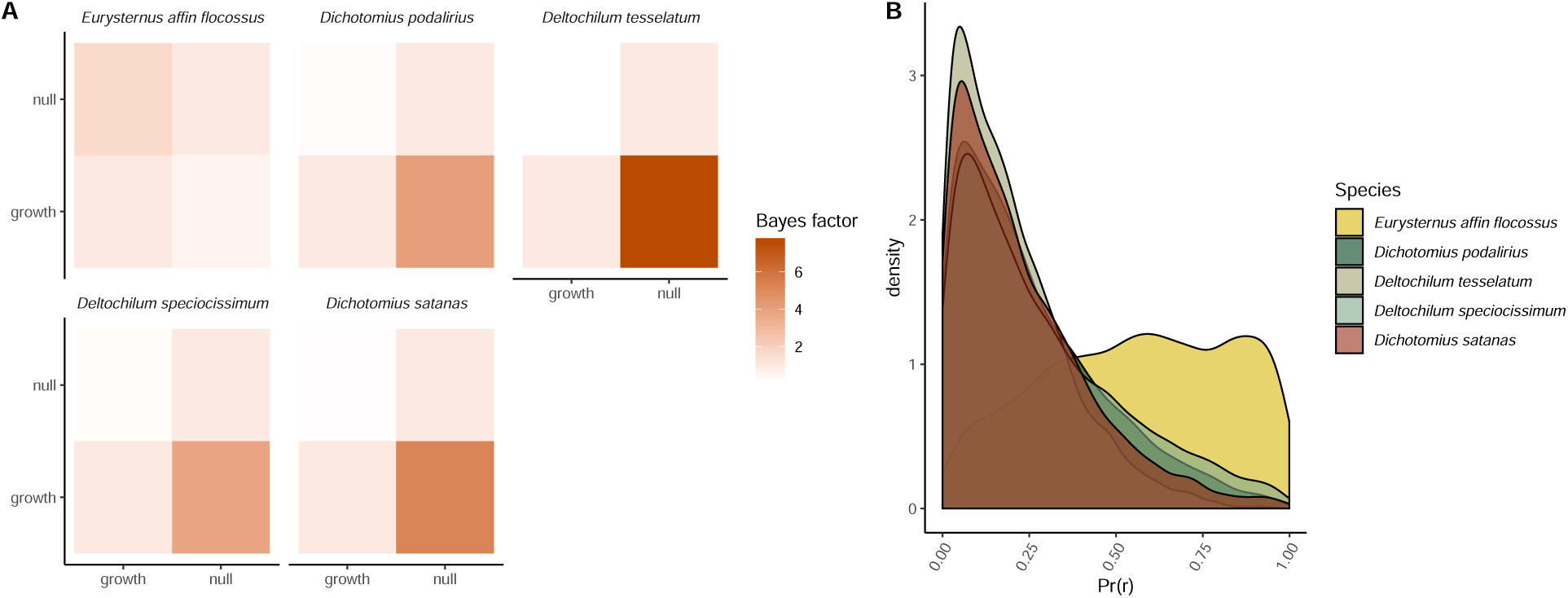
Little evidence for recent population expansion in four of five focal species. Bayes factor values for model comparisons from approximate Bayesian computation. Scores indicate the relative evidence for the model on the X axis compared to the model on the Y axis. B) Posterior probability distributions for the exponential population growth rate parameter (“r”) under the growth model from approximate Bayesian computation for all species.

### Spatial patterns of genetic diversity and Wright’s neighborhood size

Estimates of theta were uncorrelated to distance from mean sampling elevation (*χ*^2^ (1)<2.5 and p>0.1 for all species), indicating genetic diversity remains constant even in samples from near putative range limits (**Figure 5**). After correcting negative *F*_*ST*_ values and negative slope coefficients to 0 to account for artifacts of excess heterozygosity, estimates of Wright’s neighborhood size for all species were infinite (**Figure 2C**), indicating panmixia relative to the spatial scale of our sampling.

**Figure 5:**
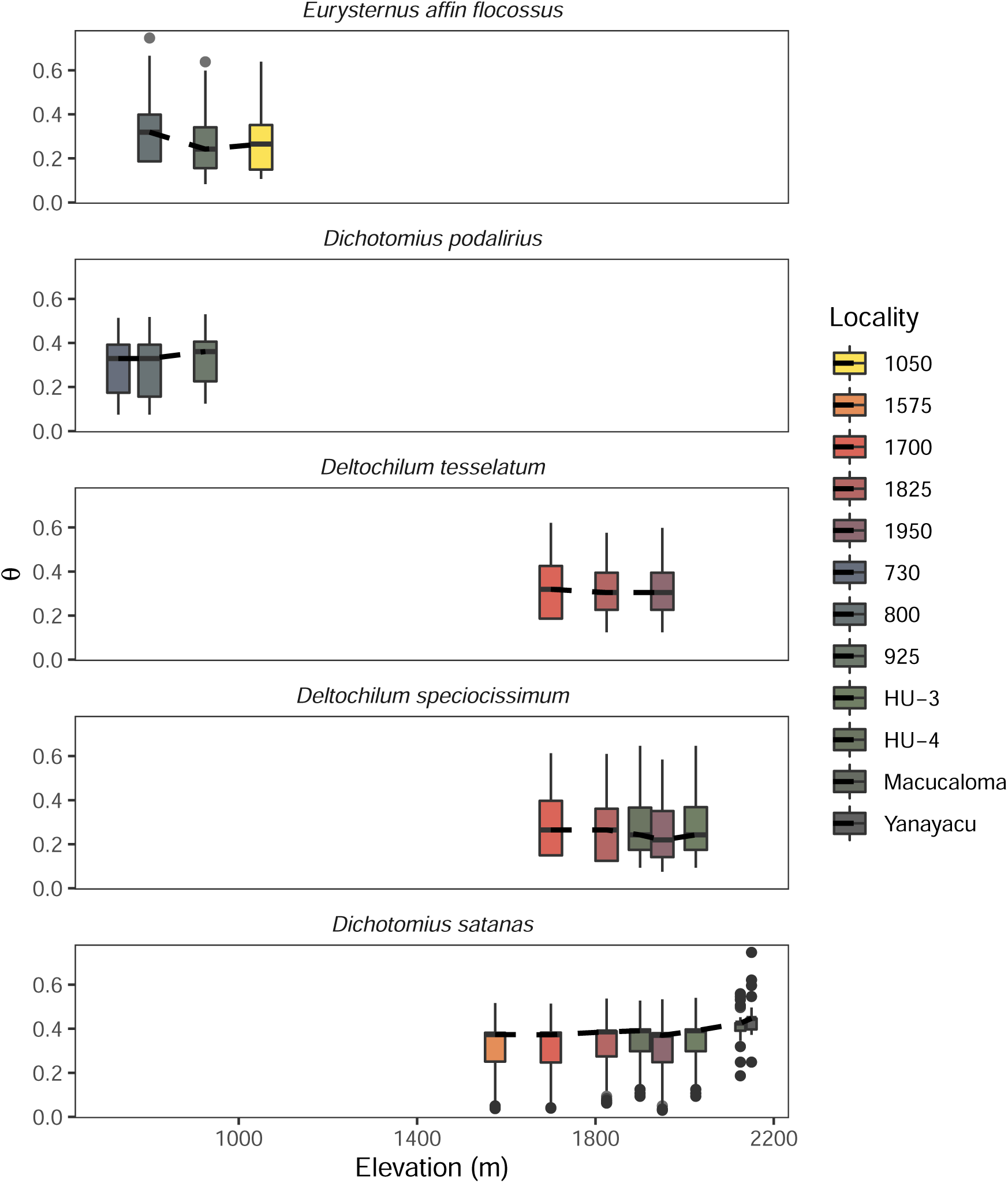
Estimates of the population-scaled nucleotide diversity (*θ*) of each sampling locality for all species. Estimates of *θ* were not correlated to distance from mean sampling elevation for any species, suggesting genetic diversity remains constant even in samples approaching the species elevational range limits.

## Discussion

Janzen’s seasonality hypothesis predicts that organisms distributed in regions with relatively limited climatic variability—as is true in much of the tropics—will evolve a reduced physiological tolerance breadth for temperature (hereafter thermal tolerance) (Janzen, 1967). In turn, narrow thermal tolerances should bias dispersal to occur most frequently between environments with similar temperature regimes, leading to neutral genetic differentiation associated with temperature and independent of geographic distance (Wang & Bradburd, 2014). Given recent evidence for the apparent ubiquity of isolation-by-environment in nature (Herrera, Medrano, & Bazaga, 2017; Manthey & Moyle, 2015; Sexton, Hangartner, & Hoffmann, 2014; Sexton et al., 2014; Shi et al., 2011; Wang & Bradburd, 2014; Weber, Bradburd, Stuart, Stutz, & Bolnick, 2017) and previous work demonstrating elevational ranges and thermal physiologies of our focal taxa broadly conform to predictions of the seasonality hypothesis (Sheldon & Tewksbury, 2014), we expected to see evidence of genetic differentiation across elevational ranges in the present study. We were therefore surprised to find no evidence of population genetic structure associated with elevation, and little evidence of isolation-by-environment in any form (Figures 2-3). Reflecting these patterns, Wright’s neighborhood sizes for all five species—proportional to the average number of potential mates for an individual given its dispersal ability—were effectively infinite relative to the geographic and environmental scale of our sampling (**Figure 2C**), indicating panmixia.

Importantly, this result appears to reflect long-term population dynamics rather than a temporary artifact of when we sampled. Over evolutionary time scales, demographic processes may obscure the signal of selection against maladaptive physiological phenotypes, biasing inferences about the drivers of genetic differentiation. For example, rapid population expansion across an environmental gradient may temporarily generate a signal of panmixia, before subsequent range contraction due to fitness costs, or the emergence of population genetic structure as a consequence of divergent selection (Gadek et al., 2018). Evidence that four of five focal species in the present study show no evidence of recent population growth (**Figure 4A**) is consistent with selection-migration equilibrium and indicates no genome-wide reduction in gene flow based on elevational origin.

What biological processes might explain rampant gene flow across elevation in Andean dung beetles? We suggest that, while initially unintuitive to us, this result could be considered a predictable consequence of their natural history. Dung beetles in Canthonini, Dichotomini and Oniticellini are strong fliers with highly developed olfactory systems (Hanski & Cambefort, 2014), and they rely on an ephemeral, randomly distributed resource for feeding and reproduction. With low mammal population density in the tropical Andes (Jiménez et al., 2010; Lizcano, Pizarro, Cavelier, & Carmona, 2002; Ríos-Uzeda, Gómez, & Wallace, 2007) and putative variation in the nutritional content of mammalian dung (Hanski & Cambefort, 2014), large, protein-rich deposits from species such as Andean bear (Tremarctos ornatus) might represent a rare “windfall” event, attracting large numbers of individuals and providing opportunities for gene flow among individuals with relatively distant locations. Given the dramatic relief of the eastern Andes and concurrently short geographic distances between upper and lower elevational range limits of the beetles, this span could easily encompass populations otherwise subject to divergent selection on thermal physiologies. Additionally, our focal species’ demonstrated ability to detect and reach dung resources (such as our traps) may make them inherently less likely to show signatures of isolation by environment.

Our findings strike a marked contrast to the results of Polato et al. (2018), to our knowledge the only other study to explicitly test Janzen’s prediction of reduced dispersal across tropical mountainsides. In their study, Polato and coauthors found gene flow across tropical elevational gradients was reduced relative to temperate elevational gradients in freshwater stream insects. We suggest the discrepancy between our conclusions may be driven by several nonexclusive mechanisms. First, as discussed above, dung beetles are highly vagile and likely show greater baseline rates of gene flow than the focal taxa in Polato et al.’s study. Second, Polato et al. did not attempt to distinguish isolation-by-environment from isolation-by-distance, perhaps due to constraints imposed by the linear nature of freshwater montane habitats. In the absence of this control, it is possible dispersal is reduced systematically across latitude, and not simply as a result of narrower thermal tolerance driven by reduced seasonality. Third, Polato et al. analyzed gene flow within morphological taxonomic units (MTUs)—putative species identified by morphology alone—rather than biological or coalescent-delimited species. They justify this by arguing reductions in gene flow between incipient species contained within a single MTU are still informative as to the validity of Janzen’s hypothesis, as speciation is one of its potential consequences. Given previous work in this system demonstrating a greater number of cryptic species at lower latitudes (Gill et al., 2016), it seems likely intraspecific reproductive isolation is a major driver of their observed reduction in gene flow across elevation. However, it is at least as plausible that cryptic speciation was driven by the evolution of reproductive isolation through processes unrelated to physiology, such as genetic incompatibilities arising during a period of allopatry prior to secondary contact and range displacement. In the absence of further evidence that estimates of gene flow in Polato et al. are truly intraspecific, it’s possible our studies examine different evolutionary time scales, and thus are not directly comparable.

Nonetheless, narrower elevational ranges in tropical compared to temperate confamilials remains consistent with a role for narrow thermal tolerance in restricting dispersal in Andean dung beetles. We hypothesize that selection against dispersal into environments with temperature regimes that exceed an organism’s thermal tolerance breadth primarily acts outside of a stable realized niche. Under this scenario, physiological tolerance breadths exceed the range of temperatures regularly experienced across the elevational distributions of our focal species, and fitness within this range shows relatively little variation. However, much below or above these elevational range limits, performance might decline rapidly and dispersers experience severe fitness costs (Hargreaves, Samis, & Eckert, 2014). More detailed characterization of performance curves across temperature in these species could inform a mechanistic niche model (Kearney & Porter, 2009) and establish whether their current distributions approach abiotic tolerance limits.

Though Janzen (1967) did not directly address the consequences of the seasonality hypothesis for speciation, our findings raise two points related to the field that merit discussion. First, as discussed above, populations in the tropics separated by climatically challenging regions should be more isolated and diverge more rapidly than similarly separated populations of temperate congeners. While an absence of robust taxonomic, phylogenetic, and distributional data have largely precluded tests of diversification rates in tropical dung beetles (but see Davis & Scholtz, 2001; Davis et al., 1999), well-designed comparative phylogeographic studies across latitude targeting common species could help evaluate this hypothesis. Second, strong constraints on thermal niche might trigger divergent selection and ecological speciation (Schluter, 2001; Nosil, 2012) in the event of niche expansion. However, the apparent stability of elevational ranges over evolutionary time scales suggests unusual circumstances might be required to facilitate this process, such as ecological release from a competitor, or a change in standing genetic variation enabling adaptation.

Our results suggest tropical dung beetles may have limited capacity to respond to climate change. Anthropogenic global warming may have a dramatic impact on tropical dung beetle communities assuming species cannot keep pace with their thermal niche through range shifts (Sheldon, Yang, & Tewksbury, 2011), though the magnitude and direction of range shifts remain unpredictable given variation in responses among closely related taxa and uncertainty in climate forecasts for tropical regions (Sheldon, 2019). Regardless, these predictions of community change ignore the potential for adaptation. Though experimental approaches to local adaptation remain the gold standard, data on the evolutionary ecology of range limits can inform predictions of the likelihood species can respond to climate changes at evolutionary timescales. Our data suggest this may be difficult for the species in the present study. In addition to evidence that most species’ ranges are at equilibrium, the absence of any decay in genetic diversity away from the elevational range center (**Figure 5**) suggests elevational ranges are not constrained by demographic effects or limited standing variation—i.e. source-sink dynamics driven by fitness costs or poor habitat quality at range limits. While it is beyond the scope of our work to identify the ultimate drivers of elevational distributions, this pattern is consistent with range limits formed by either a biotic interaction or swamping of maladaptive alleles (Sexton, McIntyre, Angert, & Rice, 2009). Assuming the latter process dominates, and niche constraints are primarily evolutionary, a complex change in patterns of gene flow might be required to permit adaptation to novel climates. We encourage future workers to explore this rich area of inquiry.

## Acknowledgements

We thank Claire Winfrey for assistance in the field, C.C. Nice at Texas State for help with library preparation, and the staff of Universidad Regional Amazónica Ikiam molecular laboratory for hosting our work. This research was conducted under permit no. MAE-DNB-CM-2017-0062 from El Ministerio de Ambiente to J. Celi, and was supported in part by an NSF IOS-1930829 to K.S.S.

## Data Accessibility

All code used in this study and an annotated digital notebook are available at https://github.com/elinck/scarab_migration. Processed data matrices are available via the Dryad Digital Repository (accession XXXX), and raw sequence read data is available at the NCBI SRA (accession XXXX).

